# MiR-4521 perturbs FOXM1-mediated DNA damage response in breast cancer

**DOI:** 10.1101/2023.02.08.527613

**Authors:** Raviprasad Kuthethur, Divya Adiga, Amoolya Kandettu, Maria Sona Jerome, Sandeep Mallya, Kamalesh Dattaram Mumbrekar, Shama Prasada Kabekkodu, Sanjiban Chakrabarty

## Abstract

Forkhead (FOX) transcription factors are involved in cell cycle control, cellular differentiation, maintenance of tissues, and aging. Mutation or aberrant expression of FOX proteins is associated with developmental disorders and cancers. FOXM1, an oncogenic transcription factor, is a promoter of cell proliferation and accelerated development of breast adenocarcinomas, squamous carcinoma of the head, neck, and cervix, and nasopharyngeal carcinoma. High FOXM1 expression is correlated with chemoresistance in patients treated with doxorubicin and Epirubicin by enhancing the DNA repair in breast cancer cells. Here, we showed that FOXM1 is a direct target of miR-4521 in breast cancer. Overexpression of miR-4521 significantly downregulated FOXM1 expression in breast cancer cells. FOXM1 regulates cell cycle progression and DNA damage response in breast cancer. We showed that miR-4521 expression leads to increased ROS levels and DNA damage in breast cancer cells. FOXM1 plays a critical role in ROS scavenging and promotes stemness which contributes to drug resistance in breast cancer. We observed that breast cancer cells stably expressing miR-4521 lead to cell cycle arrest, impaired FOXM1 mediated DNA damage response leading to increased cell death in breast cancer cells. Additionally, miR-4521-mediated FOXM1 downregulation perturbs cell proliferation, invasion, cell cycle progression, and epithelial-to-mesenchymal progression (EMT) in breast cancer. High FOXM1 expression has been associated with radio and chemoresistance contributing to poor patient survival in multiple cancers, including breast cancer. Our study showed that FOXM1 mediated DNA damage response could be targeted using miR-4521 mimics as a novel therapeutic for breast cancer.

## Introduction

FOXM1 is an oncogenic transcription factor highly expressed in rapidly proliferating cells and is absent in non-proliferating cells (Barger et al., 2019; Zhang et al., 2021). FOXM1 gene amplification and activating mutations leading to its overexpression are reported in different cancers, contributing to almost all the hallmarks of cancer (Kalathil et al., 2021). High expression of FOXM1 in breast cancer is associated with larger tumor size, lymphovascular invasion, lymph node metastasis, aggressive phenotype, resistance to chemotherapy, and an overall poor prognosis (Saba et al., 2016; Vishnubalaji and Alajez, 2021). FOXM1 promotes EMT in breast cancer by activating the SLUG gene, and metastases through interaction with SMAD3 to activate the TGF-β pathway (Xue et al., 2014). It facilitates the degradation of the ECM through the regulation of MMP2, MMP-9, uPA, uPAR, and VEGF (Saba et al., 2016). FOXM1 is a transcriptional regulator of NBS1 and facilitates double-strand break repair in breast cancer cells (Khongkow et al., 2014). Among breast cancer subtypes, a consistent overexpression of FOXM1 is seen in triple-negative breast cancer (TNBC) (Dey et al., 2020). FOXM1 expression in breast cancer is a critical regulator of MELK, an oncogenic kinase, involved in cancer cell proliferation (Bollu et al., 2020). Tumor suppressor p53 is a repressor of FOXM1 and its mutation is observed in 80% of TNBCs, leading to activation of FOXM1 expression and function in TNBC (Bollu et al., 2020). *FOXM1* overexpression has been linked to doxorubicin resistance through the regulation of XIAP and Survivin in breast cancer cells (Nestal de Moraes et al., 2015). Long noncoding RNA, TRPM2-AS mediated *FOXM1* overexpression led to development of radio resistance in gastric cancer cells (Xiao et al., 2020). *FOXM1* has been identified as a transcriptional driver of nuclear-encoded mitochondrial genes involved in mitochondrial biogenesis and mitophagy (Kondo et al., 2020). It also regulates the expression of mitochondrial antioxidant enzymes like peroxiredoxin 3 (PRX3) and acetaldehyde dehydrogenase-2 (ALDH2) (Song et al., 2015; Chen et al., 2020). *FOXM1* is an important regulator of oxidative stress and DNA damage during carcinogenesis (Park et al., 2009, 2012). Highly aggressive metastatic cancer cells show OXPHOS dependency to acquire drug resistance phenotype In OXPHOS dependent metastatic cancer cells, *FOXM1* mediated Prx3 expression facilitates suppression of ROS mediated oxidative stress induced DNA damage, senescence and apoptosis (Choi et al., 2020).

Previously, few miRNAs have been identified that regulate cancer cell proliferation, EMT, invasion, metastasis, apoptosis and drug resistance through their interaction with *FOXM1* (Liao et al., 2018). MiR-370 was reported to downregulate *FOXM1* in gastric cancer (Feng et al., 2013). MiR-134 was shown to modulate EMT by targeting *FOXM1* (Li et al., 2012). In addition, it also reverses the drug sensitivity by inhibiting *FOXM1* in lung adenocarcinoma (Li et al., 2017). During breast cancer progression, miR-671-5p, a tumor suppressor miRNA, targets FOXM1 expression and function in breast cancer cells (Tan et al., 2016). Ectopic expression of the miR-671-5p was implicated in transformation of EMT to mesenchymal-to-epithelial transition phenotype as well as an increased sensitivity to Epirubicin, 5-fluorouracil and cisplatin (Tan et al., 2016). Another study identified that miR-802 directly target *FOXM1* expression and function to inhibit the proliferation of breast cancer MCF-7 cells (Yuan and Wang, 2015). FBXL19-AS1 is a lncRNA that targets miR-876-5p in breast cancer to promote FOXM1-induced cell proliferation and inhibition of apoptosis (Dong et al., 2019, 1). Previously, miR-4521 has been identified as significantly downregulated in gastric cancer (GC) tissue specimens when compared to adjacent normal counterpart (Xing et al., 2021). It was identified that miR-4521 downregulation is linked to GC patients’ clinical stage, metastasis, and poor prognosis (Xing et al., 2021).

In this study, we identified that miR-4521 was significantly downregulated in breast cancer cell lines by performing miRNA sequencing. Further, analysis of miR-4521 expression TCGA BRCA breast cancer patient dataset showed significant downregulation of miR-4521 in tumor samples. We observed that miR-4521 was significantly downregulated in all breast cancer subtypes when compared to normal tissue. *In silico* analysis and functional assay confirmed *FOXM1* as one of the targets of miR-4521 in breast cancer cells. We observed that miR-4521 overexpressing cells have significantly high ROS levels when compared to control cells which contributes to DNA damage in breast cancer cells. miR-4521 mediated DNA damage which led to cell cycle arrest at the G1 phase. Inhibition of *FOXM1* function by miR-4521 contribute to impaired DNA repair in breast cancer cells leading to increased cell death.

Breast cancer cells with defective DNA repair capacity are vulnerable to chemotherapy which induce DNA damage and cell death. Future study exploiting miR-4521 delivery in combination with platinum drugs or PARP inhibitor in breast cancer mouse PDX model will help understand its impact as a novel therapeutic for breast cancer.

## Materials and methods

### Cell culture

Breast cancer cell lines MCF-10A (non-cancerous breast epithelial cell line) MCF-7 (luminal A), and MDA-MB-468 (TNBC) cell lines were procured from American Type Culture Collection, USA. All the cell lines were cultured in DMEM/F12 (HiMedia, India) supplemented with 10% Fetal Bovine Serum (FBS) (Gibco, USA), except MCF-10A cell lines were cultured in DMEM/F12 supplemented with 5% Horse serum (Gibco, USA), Insulin (Sigma, USA), Hydrocortisone (Sigma, USA), Epidermal Growth Factor protein (Gibco, USA). All the cell lines were grown at 37°C in a humidified chamber with 5% CO_2_ and authenticated by STR profiling.

### Small RNA sequencing

Small RNA sequencing was performed using Ion Torrent, Next Generation Sequencing platform according to standard protocols. Briefly, MCF-7, MCF-10A and MDAMB-468 cell line total RNA were isolated and after verifying the quality and quantity, small RNA enriched library was constructed using Ion Total RNA Seq Kit v2, according to the manufacturer’s instructions. Sequencing was performed using Ion PGM™ platform. Post sequencing analysis was carried out according to standard Ion small RNA Seq pipelines as per previously published protocol (Shukla et al., 2019; Kuthethur et al., 2022). *In-silico* analysis of miR-4521 target gene prediction was performed using different target gene prediction tools miRDB (Wang, 2008), TargetScan (Agarwal et al., 2015), and TarBase (Sethupathy et al., 2006). Top 10% target genes were selected from each tool and genes which were common in all 3 tools were short-listed for experimental validation.

### RNA sequencing

Briefly, RNA from empty vector control and miR-4521 over expressing MCF-7 breast cancer cells were subjected to quality and quantity verification. mRNA sequencing was performed using NovaSeq 6000 platform. Library preparation was performed using TruSeq® Stranded mRNA Library Prep (Illumina, Inc. USA) according to the manufacturer’s instructions. Sequencing was performed on NovaSeq 6000 platform using S4 paired end 200 cycle flow cell and post sequencing analysis was performed as per the standard Illumina DRAGEN RNA Seq pipelines. Pathway enrichment analysis was performed using Enrichr web-based server at Mayan cloud (https://maayanlab.cloud/Enrichr/). Briefly, differentially expressed genes from mRNA sequencing data were subjected to Reactome 2022 pathway analysis and genes were clustered into their respective pathways and ranked based on p-value.

### cDNA conversion and Real-Time PCR

cDNA conversion protocol for both miRNA specific cDNA and general cDNA was followed as described earlier (Kuthethur et al., 2022). Real-Time PCR was performed for miR-4521 expression analysis using miRNA Taqman assay (Applied Biosystems, USA) and gene expression analysis using PowerUp SYBR Green Master Mix (Applied Biosystems, USA). RNU6B was used as endogenous control for miR-4521 expression analysis, while β-Actin was used for gene expression analysis. Relative Quantity (RQ) for all the expression analysis was calculated using the formula 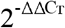.

### Plasmid Constructs

*FOXM1* 3’UTR region comprising of miRNA binding region was cloned into pmirGLO-Dual Luciferase miRNA target expression vector for miRNA luciferase reporter assay (Promega, USA) in MCF-7 cell lines. Briefly, *FOXM1* 3’UTR region was amplified from human genomic DNA, digested using plasmid specific restriction enzymes, and gel eluted along with pmirGLO empty vector (Table S1). Purified insert and vector DNA were ligated using T4 DNA ligase (NEB, USA) and transformed into DH10β chemical competent cells. Plasmid clones were isolated using alkaline lysis method and verified using restriction digestion. All the clones were confirmed by DNA sequencing before transfecting into respective cell lines.

### Transfection and Luciferase reporter assay

FOXM1 pmirGLO plasmid along with miR-4521 mimic and scrambled negative control oligo were co-transfected into MCF-7 breast cancer cell lines using Lipofectamine 3000 reagent (Invitrogen, USA), according to the manufacturer’s instructions (Invitrogen, USA). Luciferase assay was performed using Dual-Luciferase Reporter assay (Promega, USA) according to the manufacturer’s instructions using *Renilla* luciferase as internal control after 24 hours. Target gene validation was also confirmed by real-time PCR and western blotting in MCF-7 cell lines.

### Establishment of stable miR-4521 expression in breast cancer cells

For stable cell line construction, precursor miRNA sequence was amplified from human genome and cloned into pBABE-puro retroviral plasmid (Table S1). Recombinant miR-4521 retroviral titer was generated and transduced into breast cancer cells, MCF-7 and MDA-MB-468. After 48 hours of transduction, over expressing clones were selected by puromycin (1μg/ml) selection for 14 days. Individual colonies were isolated and confirmed by RT-qPCR for stable expression of miR-4521.

### Proliferation assay

8×10^4^ MCF7 cells were seeded in 35mm cell culture dishes to analyze the proliferation rate for up to 72 hours. The cells were trypsinized at indicated time points and counted using trypan blue stain and hemocytometer.

### Migration assay

pBABE control and pBABE miR-4521 cells at 90% confluency in a 6-well plate were synchronized by serum starvation for 48 hours. Wound healing assay was performed by measuring the area of scratch healed at indicated time intervals. Images were captured using Zeiss Primovert microscope equipped with Axiocam 105 camera. Images were processed and analyzed using Zen v2.3 software. The rate of cell migration and migration index were estimated as per published protocols (Xu et al., 2012).

### ROS analysis

1×10^6^ MCF-7 cells were seeded onto 60 mm cell culture dish. After 24 hours, cells were synchronized by serum starvation for 48hours. Cells were stained with 10μM concentration of DCF-DA for 30 minutes at 37°C in dark. The harvested cells were then analyzed by FACS Calibur flow cytometer and processed using Cell Quest software (BD Biosciences, USA).

### Cell cycle analysis

1×10^6^ MCF-7 cells were seeded onto 60 mm cell culture dish. After 24 hours, cells were synchronized by serum starvation for 48 hours, followed by 8 hours serum release by culturing cells in complete media. Cells were harvested, fixed with 70% ethanol at 4°C, washed with PBS and treated with 50μg/ml of RNase A for 30 minutes at 37°C. Further propidium iodide (PI) treatment was performed at 100μg/ml concentration in dark at 4°C for an hour and analyzed by FACS Calibur flow cytometer and processed using Cell Quest software (BD Biosciences, USA).

### Apoptosis analysis

1×10^6^ MCF-7 cells were seeded onto 60mm cell culture dish. After 24 hours, cells were synchronized by serum starvation for 48hours. Cells were harvested and incubated with Annexin V (Alexa Fluor 488) antibody for 30 minutes along with propidium iodide, as per the manufacturer’s instructions (Invitrogen, USA). Further cells were subjected to FACS analysis and processed using Cell Quest software (BD Biosciences, USA).

### Invasion assay

0.5% low melting agarose solution mixed with 50ng/ml EGF was spotted on 35mm cell culture dishes. 8×10^4^ MCF-7 cells with respective transfection were seeded in the same dish containing agarose spots. After respective time points, the number of cells invading the agarose spots were counted and images were acquired using Zeiss Primovert microscope equipped with Axiocam 105 camera. Images were processed and analyzed using Zen 2.3 software.

### Western blot analysis

Cells were lysed with RIPA buffer supplemented with protease inhibitor cocktails, quantified using Bradford method and 30-50μg of protein was resolved on 10-12% SDS-PAGE. Further proteins were transferred onto 0.45μm nitrocellulose membrane (Bio-Rad, USA) and blocked in room temperature with 5% BSA (HiMedia, India) solution for an hour and incubated with respective primary antibodies [FOXM1 (1:2000) (Cell Signaling Technologies, USA); active + pro Caspase-3 (1:3000) (ABclonal, Wuhan, China), B-Actin (1:5000) (Invitrogen, USA); SnaiI, CDH2, VIM (1:3000) (Cloud Clone, USA); TWIST2 (1:2000) (Abcam, USA); γH2.AX (1:2000) (Novus Biologicals, USA) and RAD51 (1:2000) (a kind gift from Dr. Ganesh Nagaraju, Indian Institute of Science, Bangalore, India) (Santa Cruz Technologies, USA)] at 4°C overnight. Blots were then incubated with secondary antibodies, either anti-rabbit IgG (1:5000) or anti-mouse IgG (1:5000) (Cell Signaling Technologies, USA) conjugated with HRP for an hour at room temperature. Finally chemiluminescent signals were developed using Clarity Western ECL substrate (Bio-Rad, USA) and visualized using ImageQuant LAS 4000 instrument (GE Healthcare, USA).

### TCGA BRCA data analysis

miR-4521 reads per million (RPM) and *FOXM1* transcripts per million (TPM) data were acquired from TCGA-BRCA datasets using TCGA Assembler tool (Wei et al., 2018) in R studio platform. Clinical data for all the TCGA breast cancer samples were downloaded from cBioPortal website. ROC curve analysis was performed using GraphPad prism tool. Survival analysis was performed using breast cancer mRNA gene chip and miRNA seq projects of Kaplan-Meier datasets.

### Statistical Analysis

Paired or unpaired Student’s t test (2-tailed) and two-way ANOVA with Turkey HSD post-hoc test was performed for respective study groups. All the experiments were performed in duplicates and data represented in the study are mean ± SD of three independent experiments. *P* value < 0.05 was considered statistically significant.

## Results

### miR-4521 is significantly downregulated in breast cancer

To identify miRNAs differentially expressed in breast cancer subtypes, we performed small RNA sequencing in breast cancer cell lines namely, MCF-7 (luminal A), MDA-MB-468 (TNBC) and compared them with non-malignant mammary epithelial cell line (MCF-10A). Differential expression analysis identified 66 miRNAs (39.1%) commonly expressed in both luminal A and TNBC subtypes, while 71 miRNAs (42%) were unique to luminal A and 32 miRNAs (18.9%) were unique to TNBC subtypes (Fig. 1A). Among the differentially expressed miRNAs, known oncogenic and tumor suppressive miRNAs such as miR-205, miR-99a, miR-200 family, miR-17, miR-21 were identified from small RNA sequencing data. Interestingly, among the top 10 differentially expressed miRNAs, miR-4521 was found to be significantly downregulated in MCF-7 and MDA-MB-468 cell lines (Fig. 1A). Validation of miR-4521 expression by qRT-PCR in breast cancer cell lines showed significant down-regulation in MCF-7 and MDA-MB-468 cell lines (Fig. 1B).

**Figure 1:**
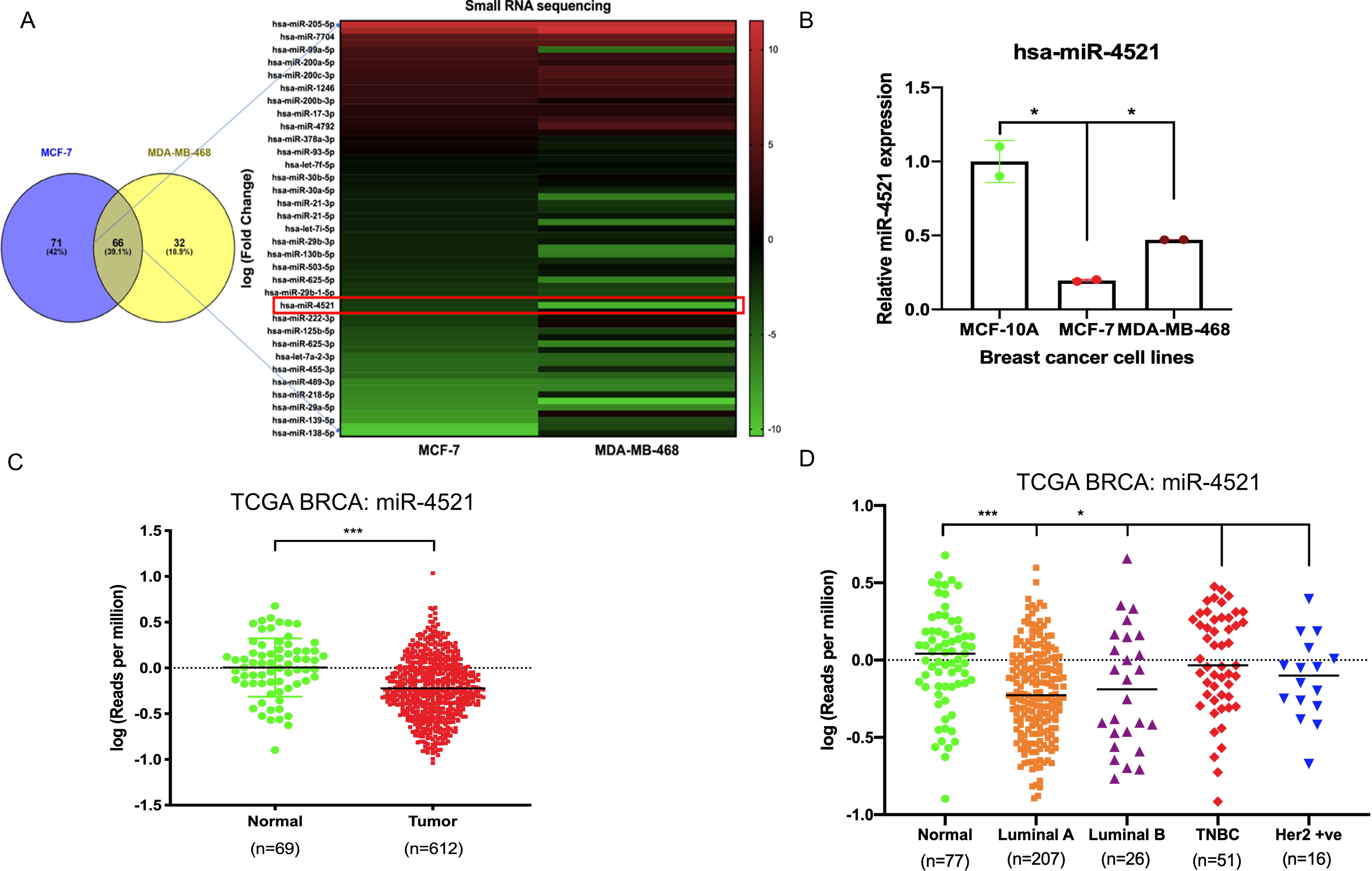
Expression of miR-4521 is significantly downregulated in breast cancer cell lines and TCGA-BRCA breast cancer patient samples. (A) miRNA sequencing identified miR-4521 as significantly downregulated in MCF-7 and MDA-MB-468 cell line when compared to MCF-10 cell line. (B) qRT-PCR expression analysis of miR-4521 in MCF-7 and MDA-MB-468 breast cancer cells compared to MCF-10A cells, RNU6B was used as endogenous control for miR-4521. Relative quantity (RQ) was calculated using the formula 2^−ddCT^; Paired Student’s t-test (two-tailed) was performed between control MCF-10A and each breast cancer cell lines, *P<0.05, (n=2). (C) TCGA-BRCA datasets from TCGA, Pan Cancer Atlas showing downregulation of miR-4521 in tumor samples when compared to normal. (D) TCGA-BRCA datasets from TCGA, Pan Cancer Atlas showing downregulation of miR-4521 in patients with different breast cancer subtypes (Luminal A, Luminal B, TNBC, and Her2 positive) when compared to normal.

Further, we systematically analyzed TCGA-BRCA Pan Cancer datasets (Cancer Genome Atlas Network, 2012) to elucidate the prognostic potential of miR-4521 in subtype, tumor stage, and nodal metastasis categories of breast cancer patient samples and observed that miR-4521 is significantly downregulated during breast cancer progression (Fig. 1C-D, S1A-B). Survival analysis using Kaplan Meier analysis showed that low miR-4521 expression is associated with poor patient survival among breast cancer patients (Fig. S1C). Similarly, ROC curve analysis of miR-4521 in TCGA breast cancer datasets showed its diagnostic potentials in different breast cancer subtypes, tumor stages, and nodal metastasis categories (Fig. S2). These combined experimental and TCGA-BRCA datasets analysis confirmed the negative association of miR-4521 with breast carcinogenesis.

### FOXM1 is a target of miR-4521

*In silico* analysis of target gene prediction using top 10% of the predicted targets from 3 different prediction tools such as miRDB (Wang, 2008), TargetScan (Agarwal et al., 2015), and TarBase (Sethupathy et al., 2006) showed *FOXM1* as the putative direct target of miR-4521 (Fig. 2A). We observed that miR-4521 binding site (MBS) on the 3’UTR region of *FOXM1* was conserved across mammals suggesting the evolutionary significance of their interaction (Fig. 2A). FOXM1 is an oncogenic transcription factor and significantly expressed in breast cancer (Park et al., 2012, 1; Sun et al., 2020, 1). Expression analysis of *FOXM1* by qRT-PCR in breast cancer cell lines confirmed *FOXM1* upregulation in MCF-7 and MDA-MB-468 cells (Fig. 2B). ROC curve analysis of *FOXM1* in TCGA-BRCA datasets showed its diagnostic potential to differentiate between normal and various subtypes of breast cancer samples (Fig. S3A). Similarly, the Kaplan-Meier survival analysis curve also showed that the high FOXM1 expression correlates with the poor survival in breast cancer patients (Fig. S3B).

**Figure 2:**
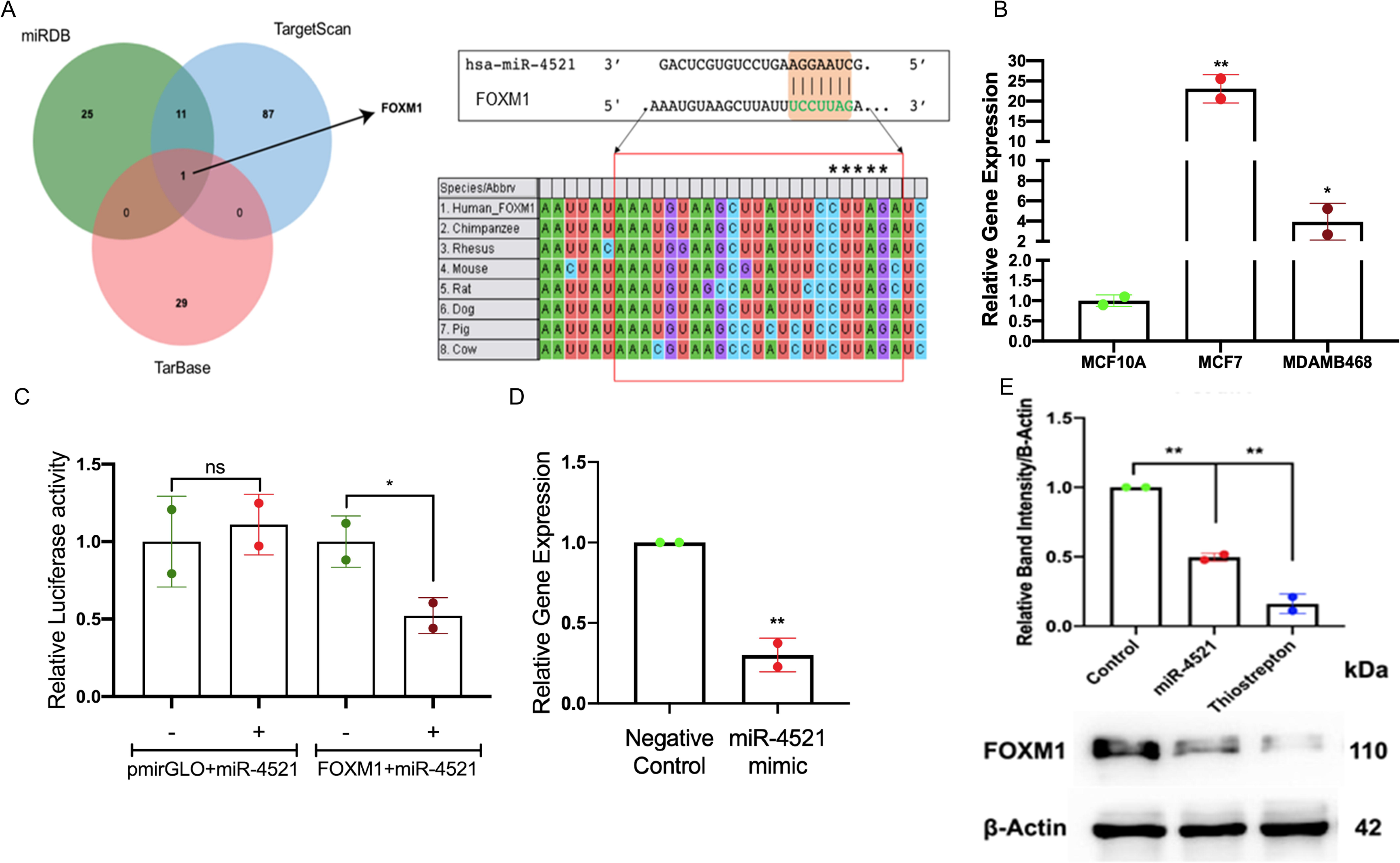
miR-4521 downregulates FOXM1 expression in breast cancer. (A) Target gene prediction for miR-4521 in three different target prediction tools: miRDB, TargetScan, and TarBase to identify putative mRNA targets. Top 10% of the targets from each tool were analyzed for common target gene and *FOXM1* was identified as target for miR-4521. Binding between miR-4521 seed region sequence with the 3’UTR of *FOXM1* mRNA sequence was conserved among mammals. (B) qRT-PCR expression analysis of *FOXM1* showed high expression in MCF-7 and MDA-MB-468 breast cancer cells compared to MCF-10A cells, β-Actin was used as endogenous control, relative quantity (RQ) was calculated using the formula 2^−ΔΔCT;^ paired Student’s t-test (two-tailed) was performed between control MCF-10A and each breast cancer cell lines, *P<0.05, **P<0.001, (n=2). (C) Dual-luciferase reporter assay upon co-transfection of 3’UTR of *FOXM1* mRNA sequence cloned pmirGLO vector along with miR4521 mimic or negative control oligos in MCF-7 cell lines after 24 hours. Relative luciferase activity was calculated upon normalizing the activity of *luciferase* enzyme to *Renilla-luciferase* enzyme activity; paired Student’s t-test (two-tailed) was performed between negative control and miR-4521 transfected groups; (+) and (-) signs indicate presence and absence of miR-4521 mimics respectively; *P<0.05, (n=3). (D) qRT-PCR analysis of *FOXM1* gene upon transfection of miR-4521 mimic for 24 hours in MCF-7 cell lines. β-Actin was used as endogenous control, relative quantity (RQ) was calculated using the formula 2^−ddCT^; paired Student’s t-test (two-tailed) was performed between negative control and miR-4521 transfected groups, **P<0.001, (n=3). (E) Western blot analysis of target FOXM1 protein abundance upon 50nM of miR4521 transfection and 1μM of Thiostrepton treatment as positive control in MCF-7 cell lines; paired Student’s t-test (two-tailed) was performed between negative control and miR-4521 transfected groups or negative control and thiostrepton treated groups, **P<0.001, (n=3).

### miR-4521 expression downregulate FOXM1 in breast cancer

Further, we validated miR-4521 and *FOXM1* interaction in breast cancer cell line MCF-7 using luciferase reporter assay. Transfection of synthetic miR-4521 mimic showed significant upregulation of miR-4521 in MCF-7 cells (Fig. S4). Luciferase reporter assay upon co-transfection of miR-4521 mimic with 3’UTR region of *FOXM1* in MCF-7 cell lines showed significant down-regulation in the luciferase activity, suggesting the direct interaction between miR-4521 and *FOXM1* (Fig. 2C). We observed decreased *FOXM1* expression in both RNA and protein levels in miR-4521 mimic transfected MCF-7 cell lines (Fig. 2D-E). Thiostrepton is a thiazole antibiotic known to downregulate FOXM1 expression and inhibit its interaction with the downstream target genes (Hegde et al., 2011). The sub-lethal concentration of thiostrepton was selected upon cytotoxicity assay in MCF-7 cell line with various concentrations ranging from 250nM to 10μM for 24 hours (Fig. S5). We observed significant downregulation in *FOXM1* protein levels of MCF-7 cell lines, upon treatment with thiostrepton (sub-lethal 1μM concentration) for 24 hours, and it is used as a positive control.

### miR-4521 mediated downregulation of FOXM1 inhibit breast cancer cell proliferation, colony formation and invasion

*FOXM1* is an oncogenic transcription factor responsible for cell cycle regulation, tumor growth, proliferation, invasion, and metastasis. Inhibition of *FOXM1* perturb cancer cell proliferation and induce apoptosis (Gartel, 2017). To understand the impact of miR-4521 overexpression in breast cancer cells, retroviral mediated stable miR-4521 expressing cells were generated using MCF-7 and MDA-MB-468 cell lines. Western blot analysis in miR-4521 overexpressing MCF-7 and MDA-MB-468 cells showed significant downregulation of FOXM1 expression (Fig. 3A). Cell proliferation assay up to 72 hours showed decreased cell proliferation rate in MCF-7 cells overexpressing miR-4521 when compared with control cells (Fig. 3B). Colony formation assay showed reduced number of colonies as well as size of the colonies in miR-4521 overexpressing cells when compared to control cells suggesting the significant reduction in tumor growth rate (Fig. 3C). Invasion assay was performed using low-melting agarose spot with 50ng/ml EGF that drives chemotactic invasion of MCF-7 cells. We observed significant decrease in invasion rate in miR-4521 overexpressing MCF-7 cells when compared to control cells (Fig. 3D). Quantification of the number of cells invading the agarose spots showed up to 80% reduction in the invasion rate compared to negative control oligo transfected cells (Fig. 3D).

**Figure 3:**
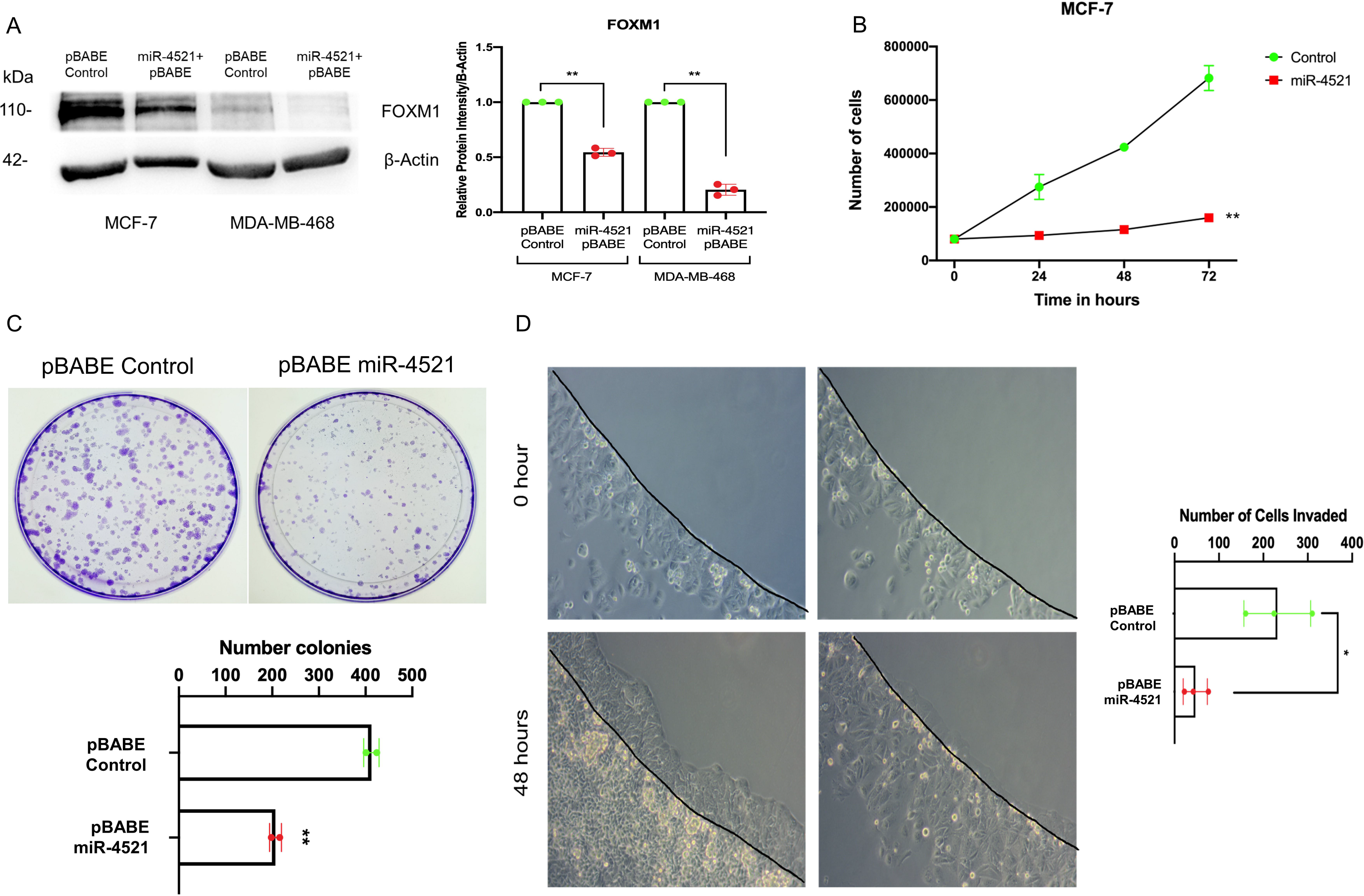
miR-4521 expression perturbs breast cancer cell proliferation, tumor growth, and invasion by downregulating FOXM1 expression. (A) Western blot analysis of FOXM1 protein levels in MCF-7 and MDA-MB-468 breast cancer cells stably expressing miR-4521. β-Actin was used as loading control; paired Student’s t-test (two-tailed) was performed between control and miR-4521 over expressing groups, **P<0.001, (n=3). (B) Proliferation rate of MCF-7 cells upon miR-4521 overexpression and control cells; paired Student’s t-test (two-tailed) was performed between control and miR-4521 transfected groups, **P<0.001, (n=3). (C) Colony formation assay among MCF-7 cells expressing miR-4521 and control cells was performed for 14 days, paired Student’s t-test (two-tailed) **P<0.001, (n=3); (D) Invasion assay upon miR-4521 over expression in MCF-7 cell lines using 0.5% low-melting agarose spots with 50ng/ml EGF for 48 hours. Quantification was performed by counting the number of agarose spot invaded cells from 3 different ROI; paired Student’s t-test (two-tailed) was performed between control and miR-4521 over expression groups, *P<0.05, (n=3).

### miR-4521 expression perturbs cell migration and mesenchymal gene expression regulated by FOXM1

RNA sequencing analysis in control and miR-4521 overexpression cell line identified downregulation of FOXM1 regulated genes involved in cell cycle (*CDC25A, CDC25B, CCNB1, CCNE1, CCND2, CDK1, PLK1, ROCK1*), DNA repair (*RFC4, PCNA, RAD51, BRCA2*), and Wnt signaling (*SKP2*) in miR-4521 overexpressing cells (Fig. 4A-B, Fig. S6, S7). *FOXM1* is a critical regulator of epithelial–mesenchymal transition (EMT) and stemness in cancer cells (Zhang et al., 2017; Nilsson et al., 2020, 1). We observed significant reduction in the protein level of the mesenchymal markers N-cadherin, vimentin; transcription factors Snai1 and Twist2 in miR-4521 overexpressing cells and *FOXM1* inhibitor thiostrepton treated MCF-7 cells when compared to control cells (Fig. 4C). Cell migration analysis by wound healing assay showed significant reduction in cell migration in miR-4521 overexpressing cells (Fig. 4D).

**Figure 4:**
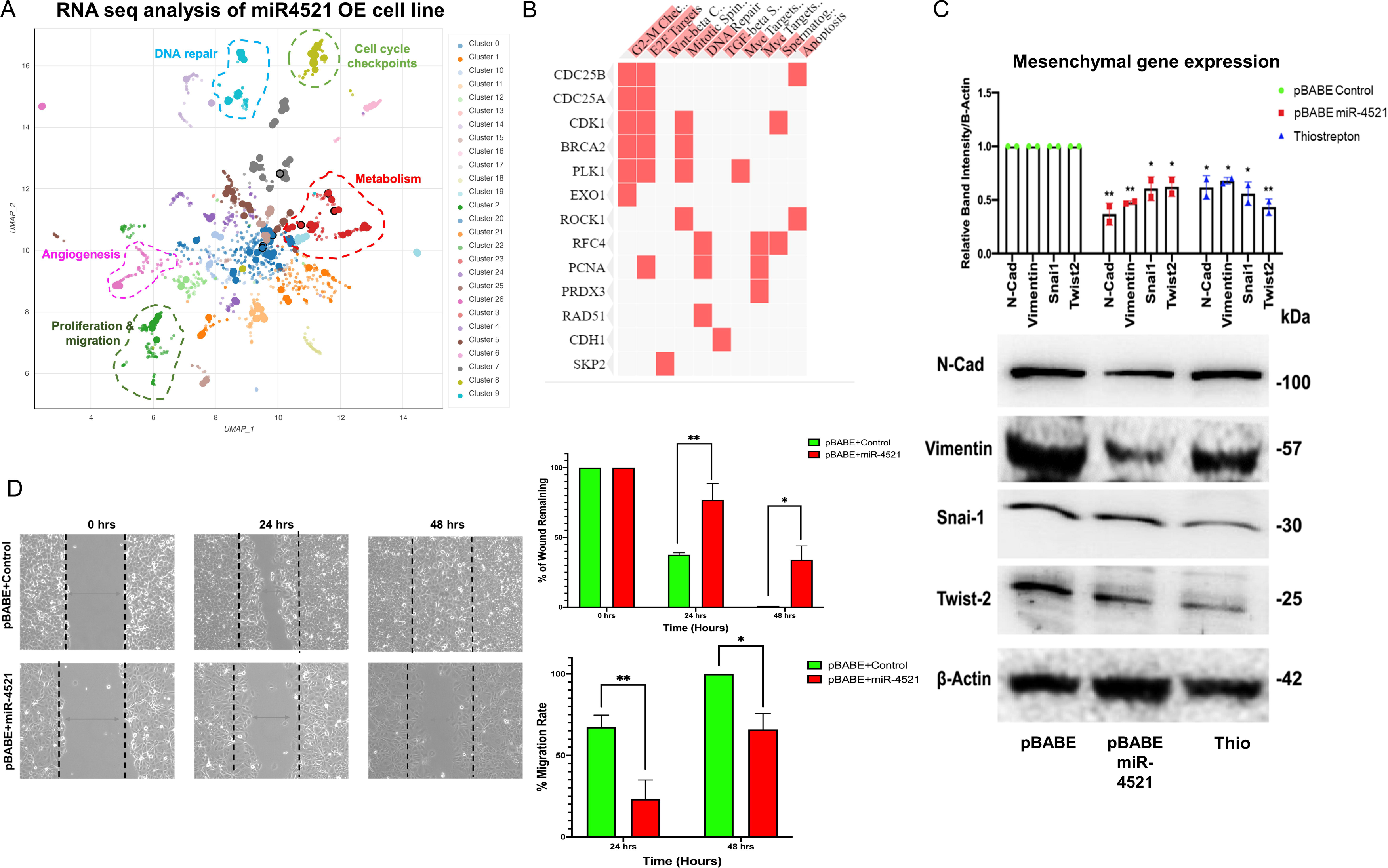
miR-4521 expression inhibits FOXM1 mediated cancer cell migration and mesenchymal gene expression in breast cancer. (A) RNA seq was performed to identify the expression pattern of FOXM1 regulated genes involved in cell cycle, replication, proliferation, invasion, migration, angiogenesis DNA repair and metabolism in miR-4521 overexpression breast cancer cells. Bokeh plot graph using Enrichr tool showing the downregulated gene enrichment pattern upon miR-4521 overexpression in breast cancer clustered based on Uniform Manifold Approximation and Projection (UMAP). (B) Cell migration was measured using wound-healing assay in breast cancer cells overexpressing miR-4521 and control cells. miR-4521 overexpression facilitate a slower migration rate in miR-4521 overexpressing cells when compared to control cells. (C) Western blot analysis of mesenchymal gene expression showed downregulation of Vimentin, Twist2, and Snail in miR-4521 overexpressing cells and FOXM1 inhibitor thiostrepton treated breast cancer cells. Densitometric analysis was performed for all the western blots upon normalizing respective test protein bands to β-Actin bands; paired Student’s t-test (two-tailed) was performed between plasmid control, pBABE-miR-4521 expressing MCF-7 cells and FOXM1 inhibitor thiostrepton treated cells, *P<0.05, **P<0.001, (n=3).

### miR-4521 induces cell cycle arrest, DNA damage and impaired DNA damage repair by downregulating FOXM1

Cell cycle analysis using propidium iodide and annexin V staining by FACS analysis showed significant increase in the percentage of G0/G1 phase cells and decrease in the S phase cells, suggesting G1 arrest in miR-4521 overexpressing cells (Fig. 5A). *FOXM1* regulates ROS induced oxidative DNA damage by transcription regulation of the expression of antioxidant genes MnSOD, catalase and PRDX3 (Park et al., 2009). Further, we analyzed the cellular ROS level in control and miR-4521 overexpressing cells and identified that miR-4521 overexpressing cells produce significantly high level of ROS when compared to control cells (Fig. 5B).

**Figure 5:**
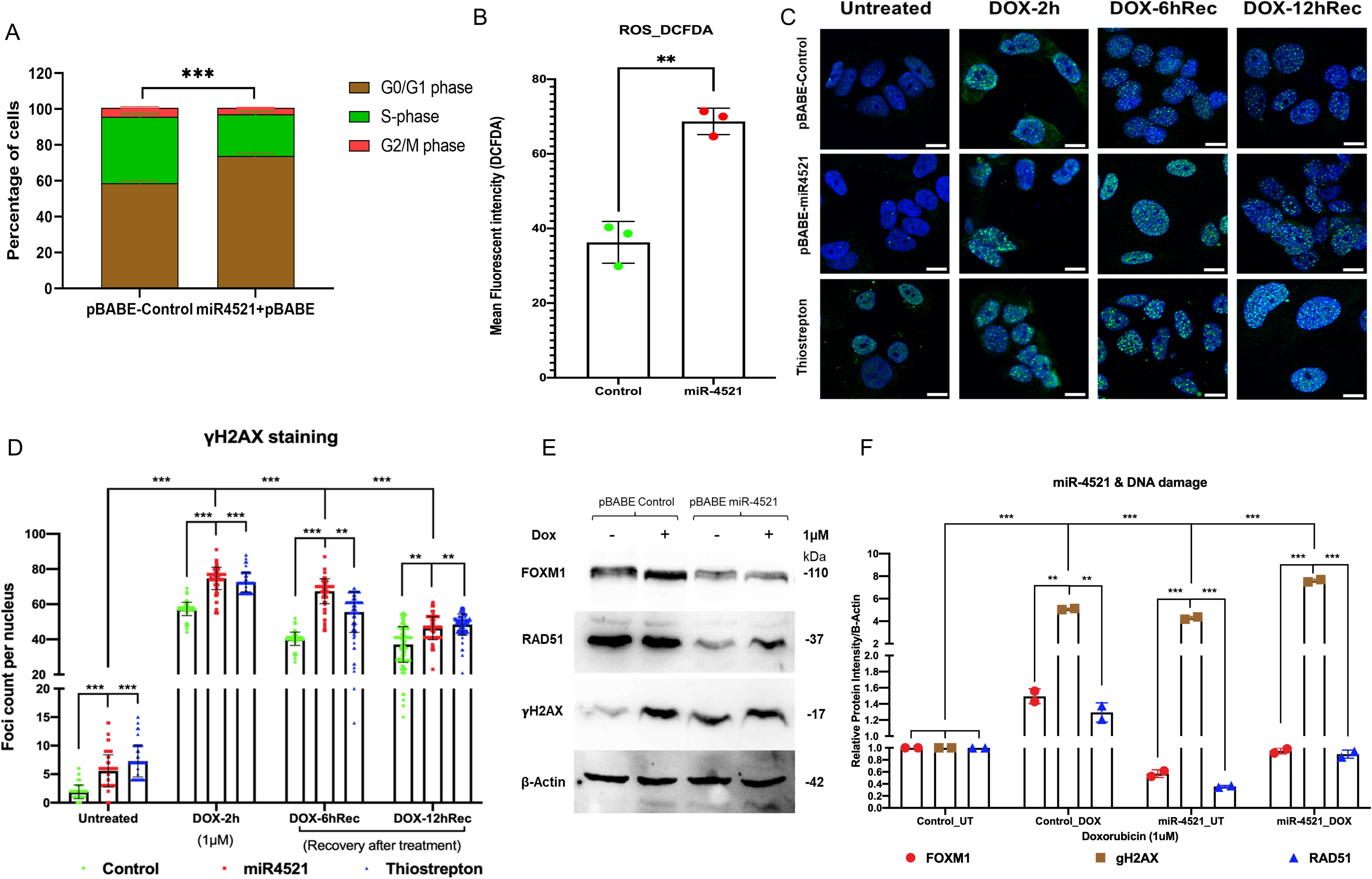
miR-4521 expression induces cell cycle arrest, increased ROS level and perturbs DNA damage response in breast cancer. (A) Cell cycle analysis of miR-4521 over expression and control cells upon synchronizing the cells at S-phase by fluorescence-activated cell sorting (FACS); paired Student’s t-test (two-tailed) was performed between control and miR-4521 over expression groups for each cell cycle stages, *P<0.05, (n=3). miR-4521 overexpression induces cell cycle arrest at G0/G1 phase by inhibiting FOXM1 mediated cell cycle regulation in breast cancer. (B) Fluorescence-activated cell sorting (FACS) analysis of cellular ROS in control cells and miR-4521 overexpressing cells using 2′, 7′-dichlorofluorescin diacetate (DCF-DA) as the molecular probe, ***P<0.0001, (n=2). Cellular ROS level was increased in miR-4521 overexpressing cells when compared to control cells contributing to cell cycle arrest and DNA damage in breast cancer cells. (C) Immunofluorescence analysis of DNA damage response in untreated and upon 1μM doxorubicin (DOX) treatment for 2 hours, followed by recovery for 6hrs and 12hrs in plasmid control cells, miR-4521 overexpressing cells, and FOXM1 inhibitor thiostrepton treated cells, (D) γH2.AX foci were quantified from 100 cells per group; two-way ANOVA with Turkey HSD post-hoc test was performed between control and miR-4521 overexpression groups or control and FOXM1 inhibitor thiostrepton treated groups at different timepoint, **P<0.001, ***P<0.0001, (n=3); Scale=25μm. (E) Western blot analysis of endogenous levels of FOXM1, RAD51, and γH2.AX in MCF-7 cell line showed increased γH2.AX expression in miR-4521 overexpressing cells suggesting DNA damage when compared to plasmid control cells. (F) Downregulation of FOXM1 showed reduced expression of RAD51, a central component of DNA double strand break repair pathway and transcriptionally regulated by FOXM1, in miR-4521 overexpressing cells. Upon induction of DNA damage by 1μM DOX treatment for 6 hours, miR-4521 overexpressing cells did not show concomitant increase in the FOXM1 and RAD51 expression when compared to control cells suggesting impaired DNA damage response in miR-4521 overexpressing cells; paired student’s t-test (two-tailed) was performed between untreated negative control and each treated or untreated groups, *P<0.05, **P<0.001, (n=3). (D) (E)

To understand the consequence of the miR-4521 mediated *FOXM1* downregulation on DNA damage response, breast cancer cells were treated with doxorubicin (DOX) at 1μM concentration for 2 hours. Upon treatment with DOX, cells were analyzed for γH2.AX foci formation in the nuclei as well as post recovery at 6 and 12 hours, in the absence of the genotoxic agent. Interestingly, both miR-4521 overexpressing and *FOXM1* inhibitor thiostrepton treated cells showed foci formation prior to DOX treatment suggesting their susceptibility to ROS induced DNA damage due to decreased *FOXM1* expression and impaired function in breast cancer cells (Fig. 5C). Upon DOX treatment for 2 hours, breast cancer cells showed increase in the damage by γH2.AX foci formation in miR-4521 expressed and FOXM1 inhibitor thiostrepton treated cells with delayed recovery rate even after 12 hours when compared to control cells (Fig. 6D). Foci quantification per 100 nuclei per group provided the accurate DNA damage response measurement as the consequence of miR-4521 mediated inhibition of *FOXM1* function in breast cancer cells. Our data confirms that miR-4521 overexpression facilitates ROS induced DNA damage with reduced DNA repair capacity by FOXM1 downregulation in breast cancer. FOXM1 is involved in DNA damage response (DDR) in cancer cells by binding to the forkhead response element region (FHRE) of DDR gene promoters and their subsequent transactivation (Zhang et al., 2012; Im et al., 2018). Western blot analysis showed increased protein levels of γH2.AX in miR-4521 overexpressing cells suggesting miR-4521 induces DNA damage in breast cancer cells (Fig. 5E). Upon DOX treatment, there was a significant increase in γH2.AX as well as FOXM1 proteins in control MCF-7 cells (Fig. 5E). However, miR-4521 overexpression cells did not show concomitant increase in FOXM1 and its downstream target gene RAD51 level upon DOX treatment suggesting impaired repair in miR-4521 overexpressing cells (Fig. 5E).

**Figure 6:**
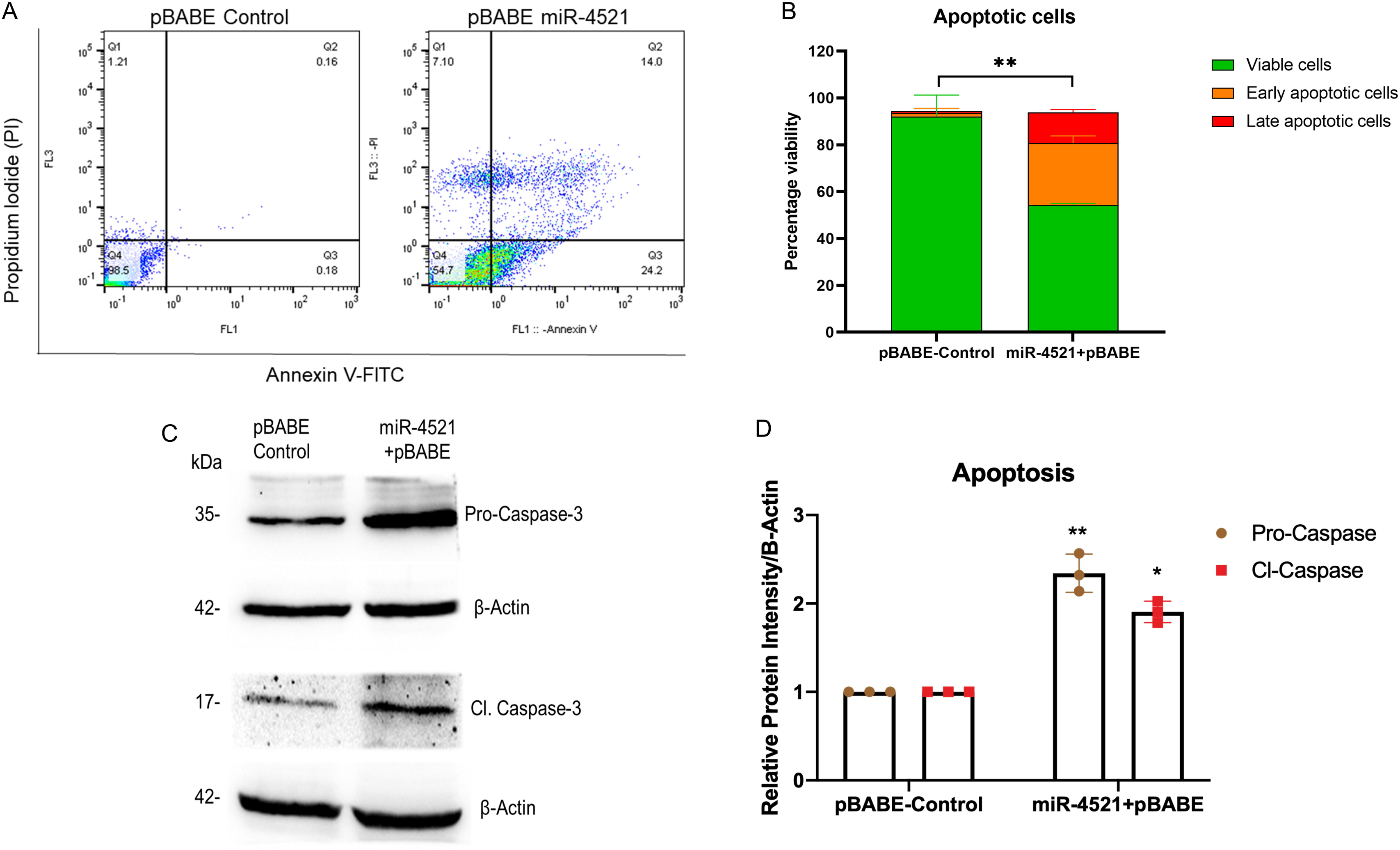
miR-4521 overexpression contribute to impaired DNA damage response and apoptosis in breast cancer cells. (A) Apoptosis analysis of Annexin V and propidium iodide (PI) stained control and miR-4521 over expressing cells. FACS analysis was performed against Annexin V-FITC and PI positive cells, where cells positive for both the stains were considered as late apoptotic cells while, cells only positive for Annexin V was considered early apoptotic cells; paired Student’s t-test (two-tailed) was performed between control and miR-4521 over expression groups for each stages of apoptotic groups; **P<0.001, (n=3). (B) miR-4521 overexpressing cells showed significant increase in early and late apoptotic cells due to increased ROS induced DNA damage, and impaired DNA damage response when compared with control cells (C) Western blot analysis of effector apoptotic protein caspase 3 showed increased cleaved caspase 3 level in miR-4521 overexpressing cells compared to control cells suggesting increased apoptosis. (D) Densitometric analysis was performed by calculating the ratio between caspase-3 and cleaved caspase-3 with respective β actin band in control and miR-4521 overexpressing cells.

### Increased miR-4521 expression contribute to DNA damage induced cell death in breast cancer

Further, we wanted to know the consequence of miR-4521 mediated DNA damage and impaired repair capacity in breast cancer cells. We performed apoptosis analysis of Annexin V and propidium iodide (PI) stained control and miR-4521 over expressing cells by FACS. We observed miR-4521 overexpressing cells showed significant increase in early and late apoptotic cells due to increased ROS induced DNA damage, and impaired DNA damage response when compared with control cells (Fig. 6A-B). We observed increased level of apoptotic protein caspase 3 and active cleaved caspase 3 in miR-4521 overexpressing cells compared to control cells further confirming increased apoptosis mediated cell death (Fig. 6C-D).

## Discussion

Breast cancer is the most common female cancer worldwide and although many patients are cured, approximately 20% develop metastasized disease, which is usually incurable. Of the different breast cancer subtypes, triple negative breast cancer (TNBC) patients, who lack expression of Estrogen or Progesterone Receptors (ER or PR) or Human EGF Receptor 2 (HER2) in their tumors, have the worst outcome and we lack a targeted therapy for this group. Among the TNBCs are patients with mutations in *BRCA1*/2, resulting in homologous recombination deficiency (HRD) and therefore impaired DNA double strand break (DSB) repair. To target this specific Achilles heel of several TNBC patients, the use of platinum-based chemotherapy and poly (ADP-ribose) polymerase inhibitors (PARPi) has been put forward as novel therapeutic approach for this subgroup. Whereas the normal tissues of the body can still cope with the DNA damage induced by these drugs, the HRD breast cancer cells cannot and die. Despite the clear benefit of these therapies, many of the patients do not respond to such treatments but experience severe side effects of chemotherapy. Moreover, in most of the patients with disseminated TNBC who do respond well initially, tumors eventually regrow and develop resistance against all available drugs. The precise resistance mechanisms are usually ill-defined, and we expect a complex network of different resistance factors.

*FOXM1*, a forkhead family of transcription factor, is involved in the regulation of cell proliferation and mitosis by regulating transcription of p27kip1, cyclin D1, cdc25, KIF20A, and CENP-A in breast cancer (Wonsey and Follettie, 2005). Depletion of *FOXM1* in cancer cell leads to chromosomal instability (Wonsey and Follettie, 2005) and regulate various oncogenic pathways leading to tumor proliferation, migration, invasion as well as DNA damage repair (Zona et al., 2014; Gartel, 2017). Hence, understanding the molecular mechanisms of *FOXM1* regulation could shed light on new avenues for breast cancer diagnosis and therapeutics. Previously, *FOXM1* was identified as one of the targets of miR-4521 in medulloblastoma (18) and gastric cancer (17). We identified that miR-4521 was significantly downregulated in breast cancer cell lines and breast cancer patient tissue samples (TCGA-BRCA datasets). Further, higher expression of miR-4521 was also associated with the better survival of breast cancer patients and emphasizes its prognostic and diagnostic potentials in breast cancer progression. Similarly, downregulation of miR-4521 expression was reported in various types of cancers such as, medulloblastoma, renal carcinoma, gastric cancer, and hepatocarcinoma (Feng et al., 2019, 129; Senfter et al., 2019; Xing et al., 2021). Being a oncogenic driver, *FOXM1* is ubiquitously expressed in rapidly proliferating and immortalized cells, while it is absent in differentiated cells (Li et al., 2009). Its overexpression is observed in various cancers like hepatocellular carcinoma, lung adenocarcinoma, hypopharyngeal squamous cell carcinoma, esophageal carcinoma and breast carcinoma where it is associated with an overall worse prognosis (Li et al., 2009; Park et al., 2012; Wei et al., 2015). It has also been implicated in the migration and invasion of cancer cells via the regulation of *MMP-2* and *MMP-9* in several cancers, *PTTG1* in colorectal cancer, and *YAP1* in triple negative breast cancer (Wang et al., 2008, 1; Zheng et al., 2015, 1; Sun et al., 2020). miR-4521 being the post-transcriptional regulator of *FOXM1*, it significantly reduces FOXM1 expression in both RNA and protein levels. Overexpression of miR-4521 showed decreased cell proliferation, invasion and altered cell cycle profile in breast cancer cells. *FOXM1* overexpression is also involved in the promotion of epithelial-to-mesenchymal transition and cancer metastasis. It has been reported to upregulate EMT-transcription factors like Snai1 and Snai2 in breast and lung adenocarcinoma (Park et al., 2012; Wei et al., 2015). We observed decreased expression of mesenchymal markers in miR-4521 overexpressing cells due to the reduced expression of FOXM1.

*FOXM1* expression is also associated with the development of chemoresistance to DNA-damaging agents like doxorubicin and 5-fluorouracil, and taxanes like docetaxel and paclitaxel (Park et al., 2012; Xie et al., 2017; Huang et al., 2019). The modulation of drug resistance occurs through the FOXM1-dependent regulation of antiapoptotic factors like Survivin and XIAP; ATP binding cassette proteins like ABCC10, ABCG2 and ABCA2; and epigenetic regulator UHRF1 (Nestal de Moraes et al., 2015; Xie et al., 2017; Yuan et al., 2018; Huang et al., 2019, 2). In addition, FOXM1 is an upstream regulator of various DNA damage repair genes such as *XRCC1*, *BRCA2*, *RAD51*, *BRIP1* and *NBS1*, thereby enabling the tumor cells to resist the action of the genotoxic agents (Zona et al., 2014). Overexpression of miR-4521 leads to increased DNA damage in breast cancer cells possibly due to impaired repair capacity of breast cancer cells. Increased FOXM1 expression is critical for reducing ROS levels in proliferating cells thereby controlling ROS mediated oxidative DNA damage in cancer cells (Park et al., 2009). Inhibition of FOXM1 leads to increased ROS induced DNA damage and apoptotic cell death in cancer (Halasi and Gartel, 2012; Halasi et al., 2013). Overexpression of miR-4521 in breast cancer cell led to ROS induced DNA damage and increased cell death suggesting its potential as novel FOXM1 inhibitor which can be further exploited in breast cancer mouse model alone or in combination with platinum drugs or PARP inhibitor (Fig. 7).

**Figure 7:**
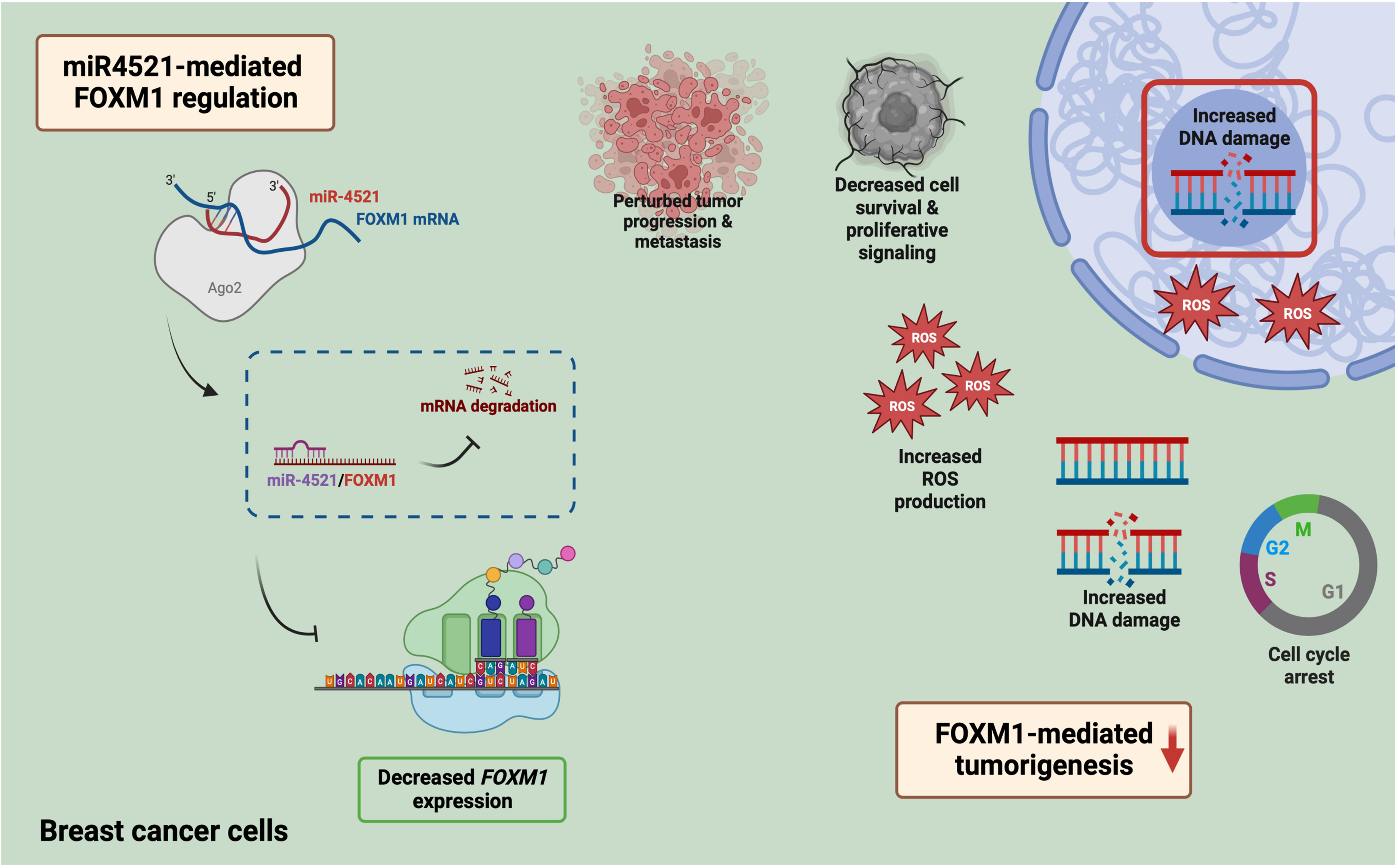
Mechanism of miR-4521 mediated inhibition of FOXM1 expression and perturb DNA damage response in breast cancer. Overexpression of miR-4521 perturbs cell cycle progression, invasion, and migration in breast cancer cells. miR-4521 mediated FOXM1 downregulation perturbs DNA damage response in breast cancer cells and increase apoptosis leading cell death.

## Conclusion

We have conclusively shown that miR-4521 perturbs *FOXM1* function and promote significant reduction in cancer cell proliferation, cell cycle arrest at G1 phase, decreased cell migration and reduced expression of mesenchymal marker expression in breast cancer. Our study has identified that miR-4521 overexpression perturbs *FOXM1* mediated transcription regulation of cell cycle progression contributing to G1 arrest and ROS mediated DNA damage in breast cancer cells. Further, miR-4521 mediated *FOXM1* downregulation impairs DNA damage response in breast cancer cells leading to increased cell death in breast cancer. We have developed mouse xenograft model to study the impact of miR-4521 overexpression on breast tumor growth and future study will explore the therapeutic role of miR-4521 alone or in combination with radiation and platinum-based chemotherapy in breast cancer.

## Abbreviations

DMEM/F12: Dulbecco’s modified eagle medium and Ham’s F-12 medium
EGF: Epidermal growth factor
EMT: Epithelial to mesenchymal transition
FACS: Fluorescence-activated cell sorting
FBS: Fetal Bovine Serum
FHRE: Forkhead response element region
HRP: Horseradish peroxidase
MBS: MicroRNA Binding Site
RISC: RNA induced silencing complex
ROC curve: Receiver operating characteristic curve
ROS: Reactive oxygen species
TNBC: Triple negative breast cancer

## Declarations

**Ethics approval and consent to participate:** Not applicable

**Availability of data and material:** The data that support the findings of this study are included in the manuscript file.

**Supporting information:** This article contains supporting information.

**Consent for publication:** We declare that all named authors have read the manuscript and have given consent to publish.

**Competing interests:** The authors declare no competing interest.

**Funding:** This work was supported by Science and Engineering Research Board (SERB), Department of Science and Technology, Government of India (YSS/2015/001051, and CRG/2020/004681). RK was supported by Indian Council of Medical Research-Senior Research Fellowship (File No: 2019-0278/GEN-BMS), Government of India.

## Authorship contribution

**Raviprasad Kuthethur**: Formal analysis, Investigation, Validation, Data Curation, Writing-Original draft preparation. **Divya Adiga**: Formal analysis, Investigation, Validation. **Amoolya Kandettu:** Formal analysis, Investigation, Validation. **Maria Sona Jerome**: Formal analysis, Investigation, Data Curation. **Sandeep Mallya**: Formal analysis, Data Curation, Validation. **Kamalesh Dattaram Mumbrekar**: Resources, Writing –Review & Editing. **Shama Prasada Kabekkodu**: Resources, Writing –Review & Editing. **Sanjiban Chakrabarty**: Conceptualization, Methodology, Validation, Formal analysis, Investigation, Supervision, Writing- Original draft preparation, Writing –Review & Editing, Funding acquisition.

## Acknowledgements

This work was supported by Science and Engineering Research Board (SERB), Department of Science and Technology, Government of India (YSS/2015/001051, and CRG/2020/004681). RK was supported by ICMR-SRF fellowship (File No: 2019-0278/GEN-BMS), Government of India. The infrastructure support and funding from DST-FIST, Government of India, TIFAC-CORE, DBT-BUILDER grant, Government of India, VGST Karnataka, K-FIST and Manipal Academy of Higher Education is gratefully acknowledged.

