## Supplementary file for "MiR-4521 perturbs FOXM1-mediated DNA damage response in breast cancer"

**Running Title**: miR-4521 perturbs FOXM1 function in breast cancer

Supporting information:

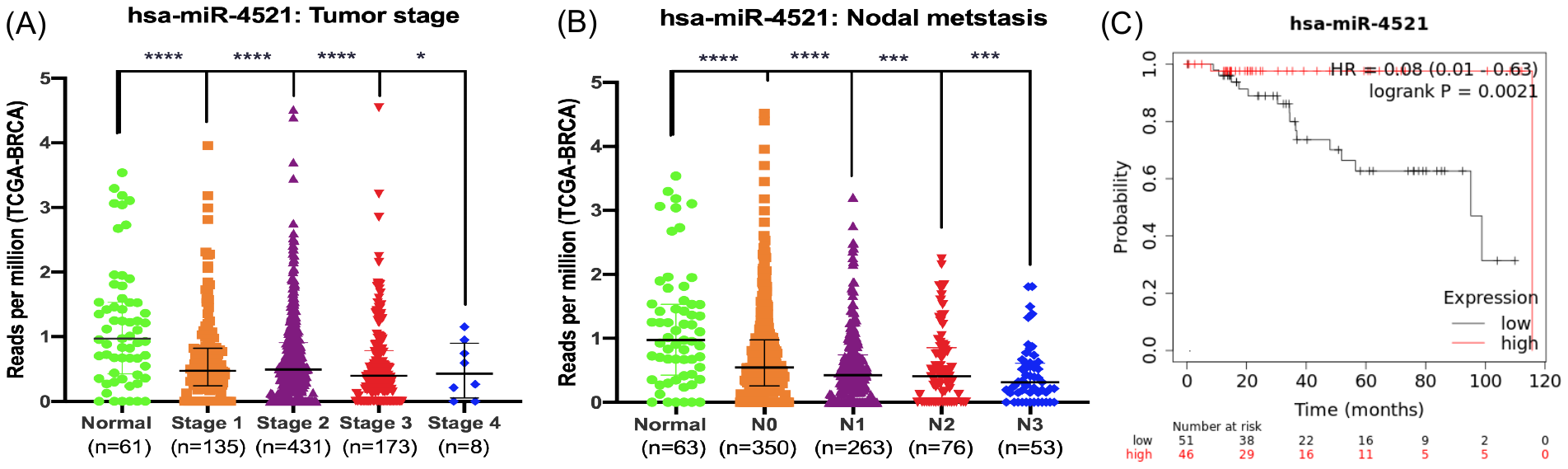

**Fig S1: Prognostic potentials and survival analysis of miR-4521 and breast cancer:** Expression analysis of miR-4521 in TCGA-BRCA patient samples through (A) tumor stages and (B) Nodal metastasis categories. (C) Survival analysis of miR-4521 in breast cancer samples through Kaplan-Meier plotter.

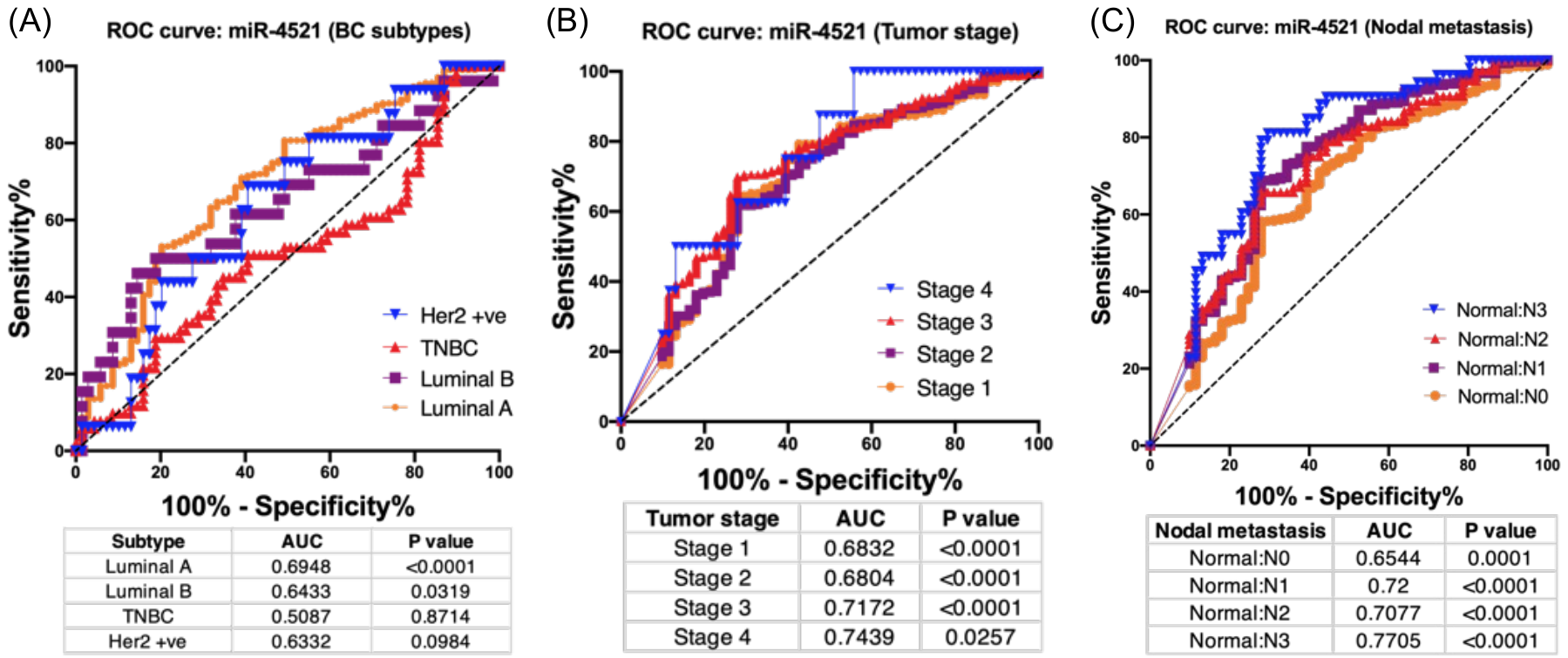

**Fig S2: Diagnostic potentials of miR-4521 and breast cancer:** Diagnostic potentials of miR-4521 in TCGA-BRCA patient samples through ROC curve analysis: (A) breast cancer subtypes, (B) tumor stage, and (C) nodal metastasis categories.

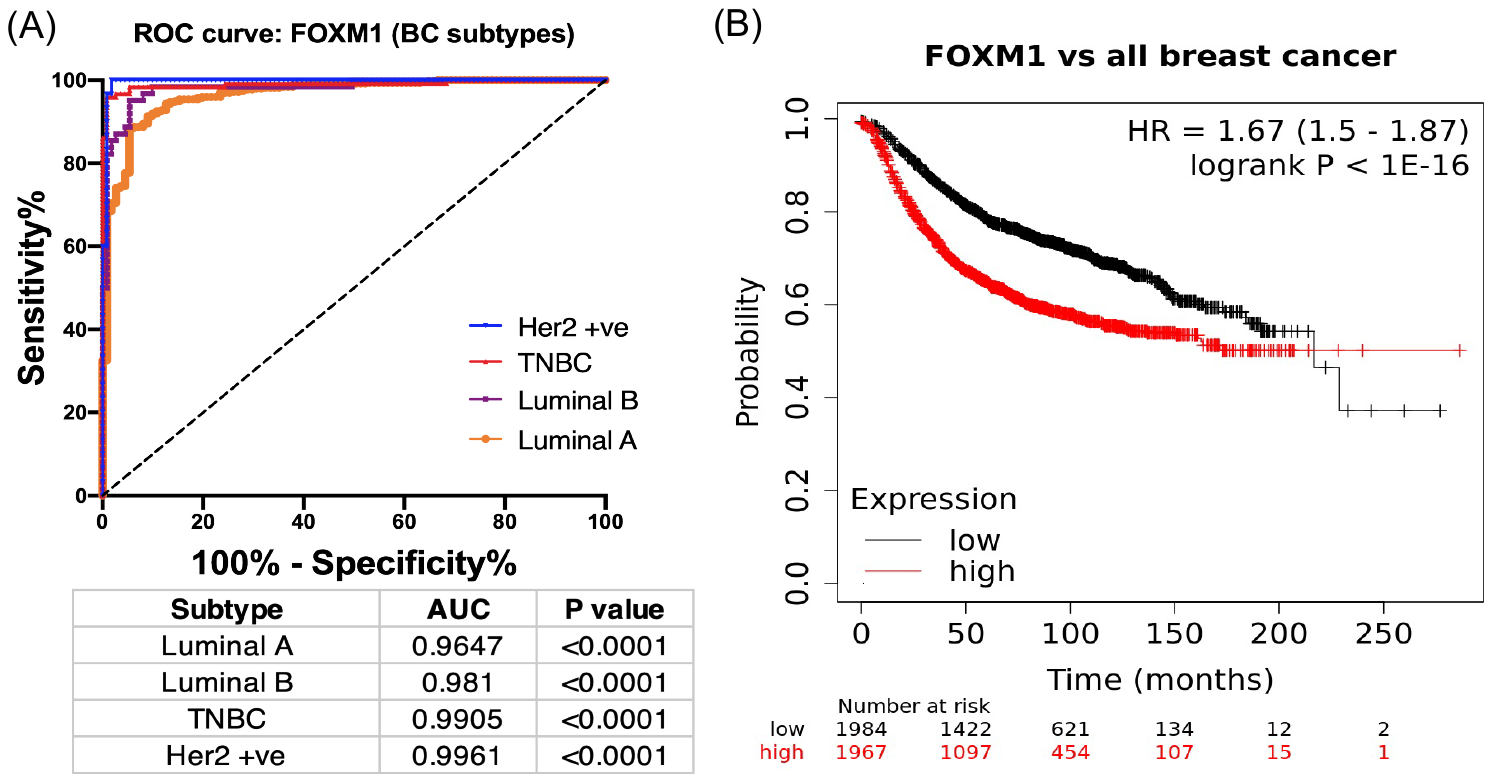

**Fig S3: Diagnostic potential and survival analysis of *FOXM1* and breast cancer:** (A) Diagnostic potentials of *FOXM1* in TCGA-BRCA patient samples through ROC curve analysis of breast cancer subtypes. Kaplan-Meier survival analysis curve depicting the high survival probability of breast cancer patients with low FOXM1 expression.

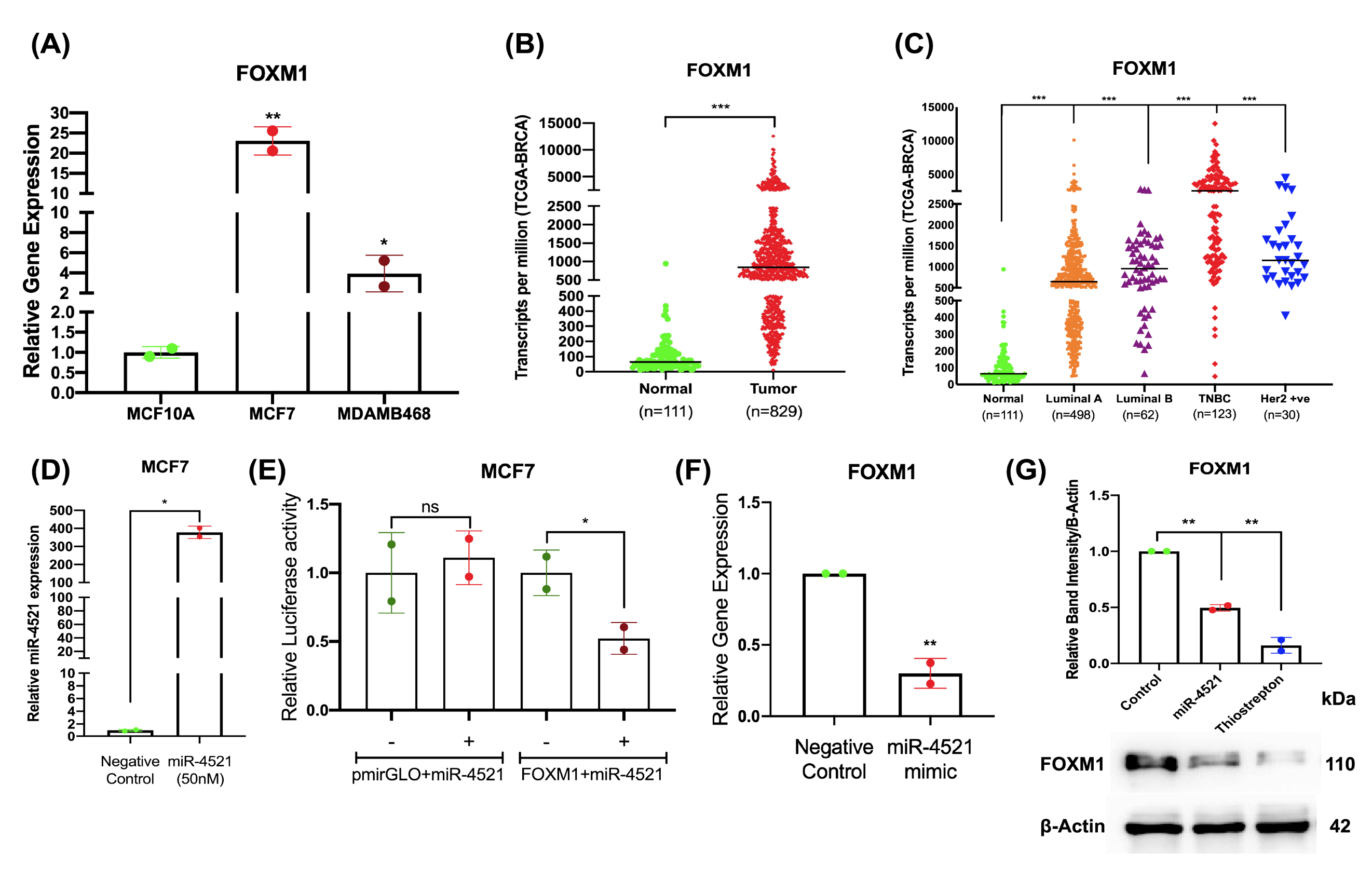

**Fig S4**: qRT-PCR analysis to measure the abundance of miR-4521 upon transfection of 50nM of miR-4521 mimic for 24 hours in MCF-7 cell lines; paired Student’s t-test (two-tailed) was performed between negative control and miR-4521 transfected MCF-7 cell line, *P<0.05, (n=2).

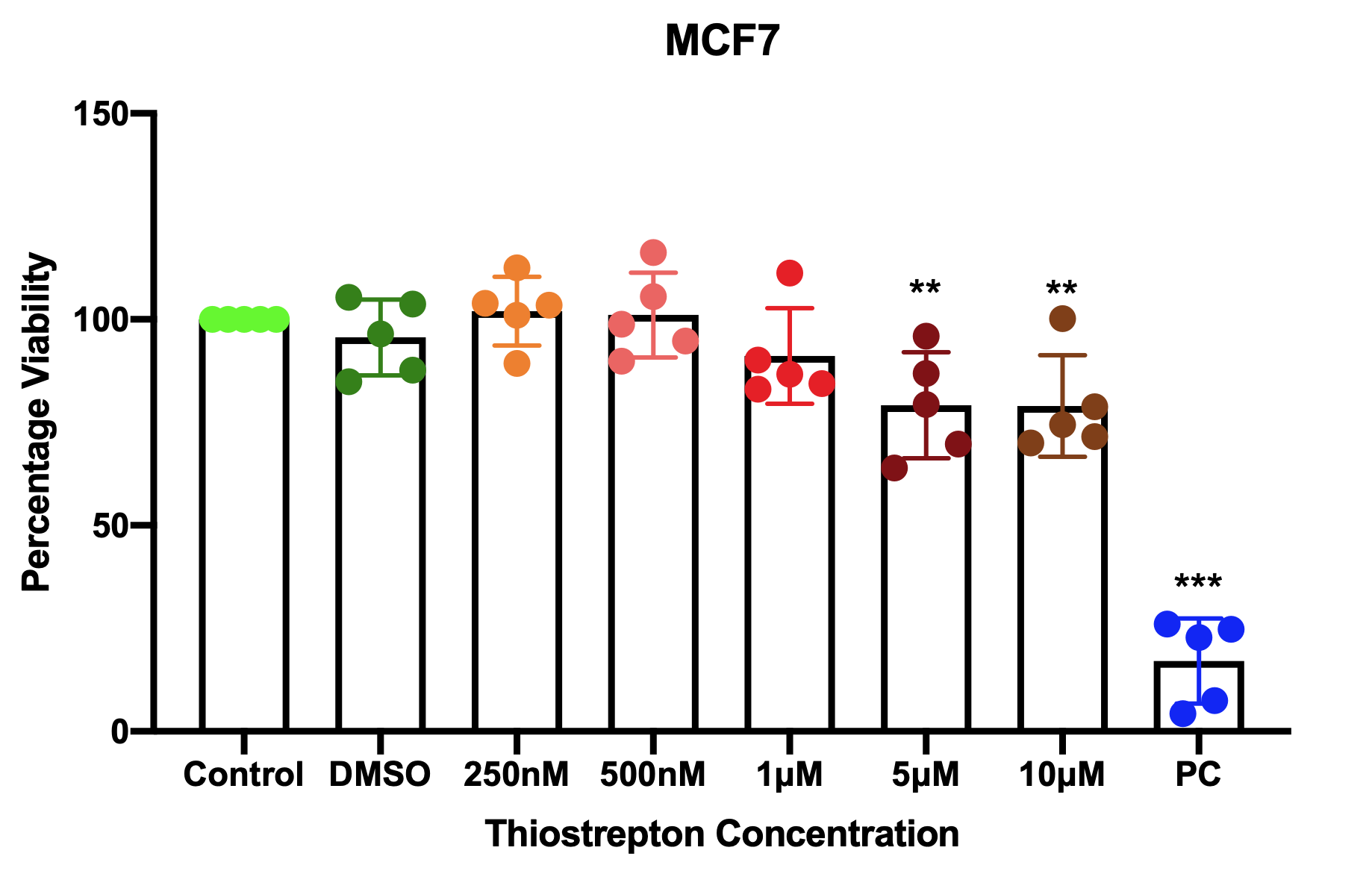

**Fig S5: Cytotoxicity levels of Thiostrepton.** Breast cancer cell line MCF-7 was treated with respective concentrations of thiostrepton for 24 hours along with DMSO as vehicle control and 5μg/ml of mitomycin as a positive control.

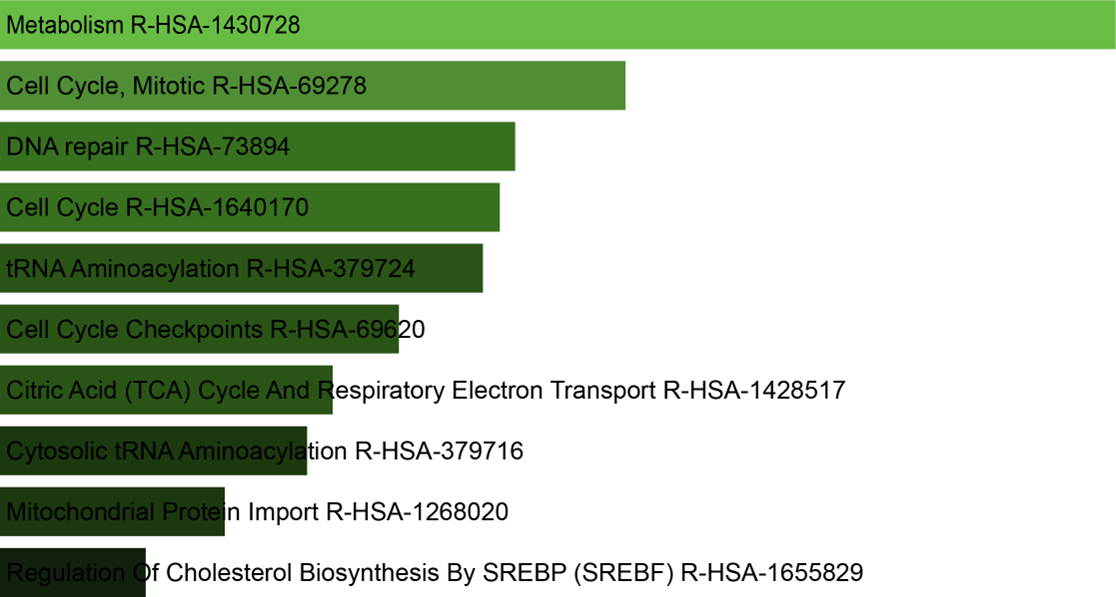

**Fig S6:** Pathway enrichment analysis of downregulated genes identified in miR-4521 overexpressing breast cancer cells by RNA sequencing using Enrichr tool. Downregulated genes belonging to respective pathways were clustered based on their p-value ranking (P<0.05).

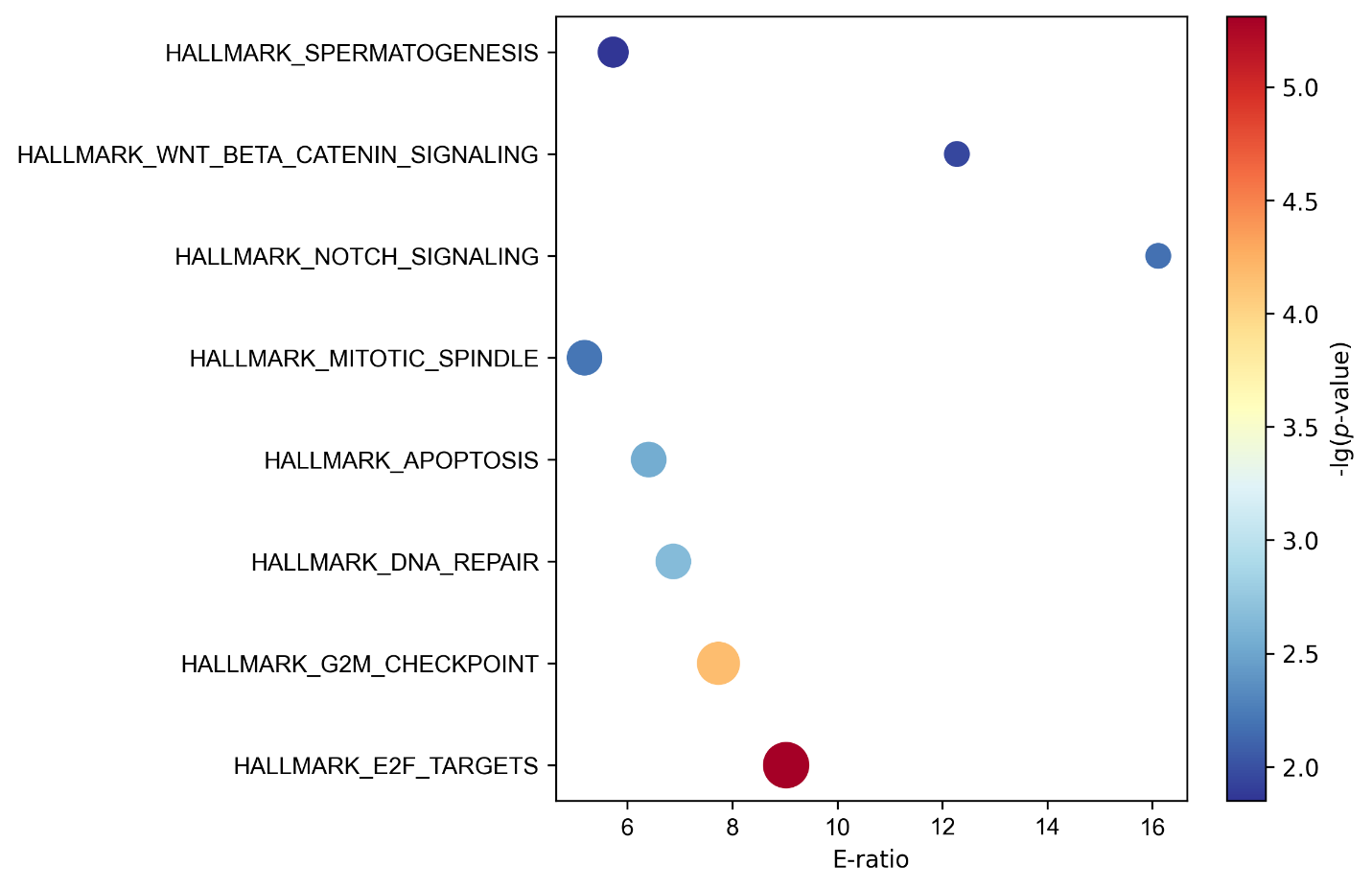

**Fig S7:** Analysis of hallmark gene expression showed significant downregulation in G2M checkpoint, E2F targets, DNA repair, Wnt signaling, Notch signaling, and mitotic spindle formation in miR-4521 overexpressing cells.

**Table S1**: List of primers used in the study.

| Sl. No. | Name | Sequence | Primer Length | Restriction Enzyme |
| --- | --- | --- | --- | --- |
| 1 | FOXM1-Cloning-For | CAGCGAGCTCCTGTTTCCATTCTCTG | 26 | SacI |
| 2 | FOXM1-Cloning-Rev | GTTAGTCGACGTGTTCTCAAGCTGGC | 26 | SalI |
| 3 | FOXM1-GE-For | GATGAGTTCTGATGGACTG | 19 | NA |
| 4 | FOXM1-GE-Rev | CATGTAAGAGTAGGGTGG | 18 | NA |
| 5 | miR-4521 cloning-F | GGGTTCGAATCCCATCCT | 18 | NA |
| 6 | miR-4521 cloning-R | ACTGAATGGAGTGACTGG | 18 | NA |
